# Highly under-actuated dynamic manipulation: Dice stacking is mostly open-loop

**DOI:** 10.64898/2026.01.08.698399

**Authors:** Noah I. Eckstein, Matthew Lerner, Manoj Srinivasan

## Abstract

Humans’ ability to grasp and dynamically manipulate objects with their hands is unmatched by current robots. To better understand human dynamic manipulation, we studied dice stacking, a task in which humans form a vertical stack of dice from a set of initially unstacked playing dice using an overturned cup and the surface of a table. This task is high dimensional and under-actuated, so it may superficially seem an incredible feat of state estimation and feedback control, but we show that this task is amenable to open-loop strategies. We simulated a cup with dice oscillated by fixed arm movement patterns using two different computer simulation frameworks with different contact models. These simulations showed that, for a range of arm and wrist movements, the dice naturally stack without any dice state feedback. We verified the predictions of these simulations with a physical robot. Thus, we have added dice stacking to the small list of dynamic manipulation tasks that can be robustly performed open-loop. We speculate that, for highly under-actuated tasks, humans may be biased to learn open-loop strategies over state feedback strategies. Future work could investigate the presence of such a bias in humans and its potential value for reinforcement learning algorithms.

## 1 Introduction

Humans can perform remarkably sophisticated dynamic object manipulation with our hands, but replicating this in robots remains difficult despite progress in deep reinforcement learning [1] and imitation learning [2]. Bridging this gap requires understanding the principles underlying human dynamic manipulation, especially in tasks humans learn easily. Here, we study one such task, dice stacking, which many people learn in minutes, despite seeming like a sleight-of-hand magic trick.

Dice stacking [3] involves using a cup and table to build a vertical stack of dice (Figure 1a). Online videos show its impressive variants, but we focus on a simple one where all dice start beneath the cup and are stacked solely through cup motion (Supplementary Video V1). The task seems impossible: the cup moves all dice simultaneously, providing no independent control. Each die has six degrees of freedom, so the system has at least 6*N*_dice_ unactuated degrees of freedom.

**Figure 1:**
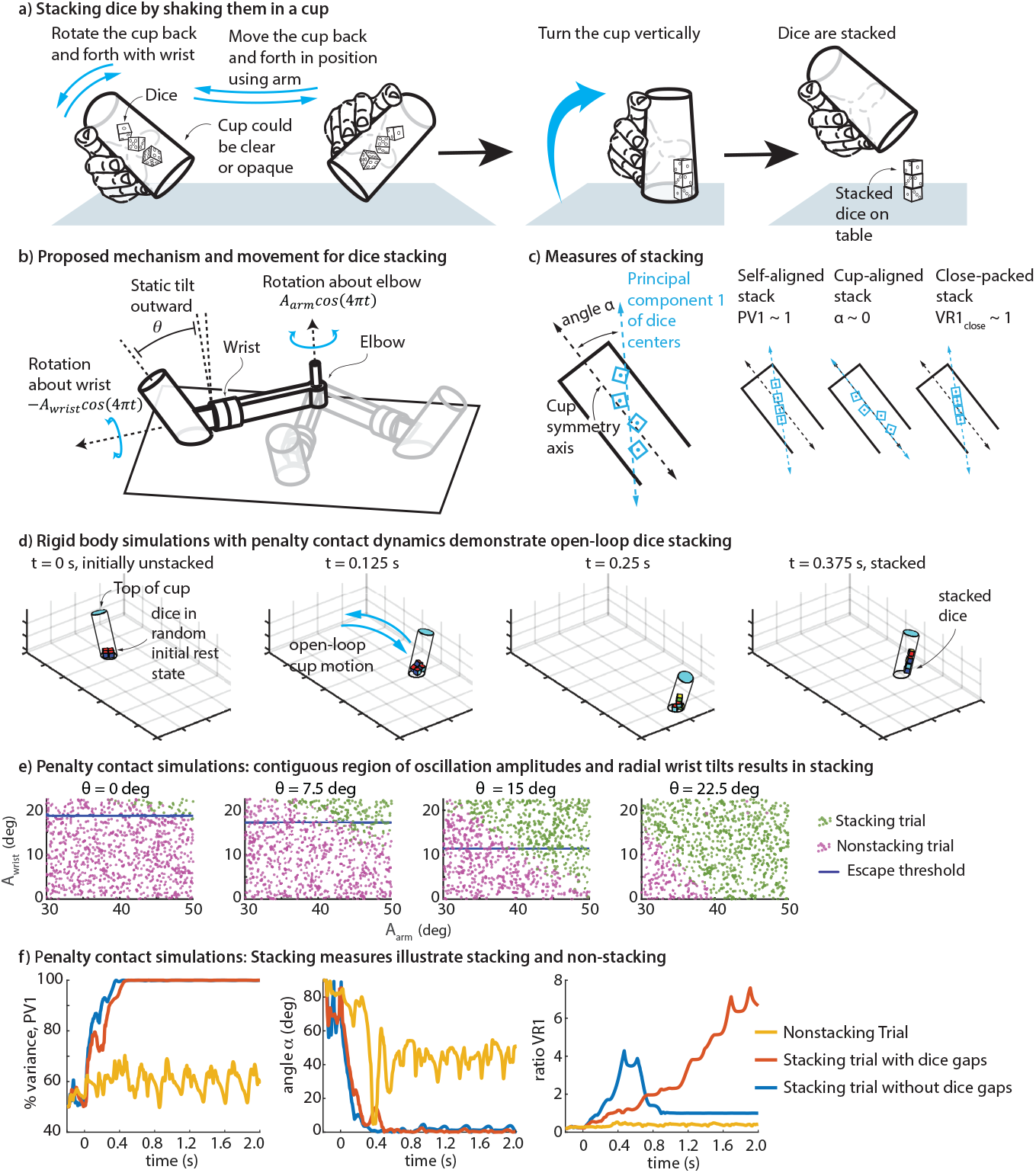
Dice stacking and performing it with open-loop arm motions. a) Dice stacking. This study focuses on the first step of dice stacking, where the dice are shaken in the cup until they form a stack, showing that it is likely open-loop stable. b) Simplified arm model with wrist pronation/supination and shoulder rotation; radial cup tilt *θ* is constant during each trial. c) Three stacking measures: *α* (angle between the first principal component of the dice center of mass distribution and the cup axis), PV1 (percent variance captured by the first principal component of dice centers), and VR1_close_ (measure of whether the dice flush against each other while stacked). d) Computer simulations with a penalty contact model show dice stacking with open-loop arm and hand motions. Dice go from randomly arranged to being in a line and flush against the cup wall in about 0.375 seconds, 3/4 of an oscillation. e) Oscillation amplitudes and radial wrist tilt yielding stacking in penalty contact simulations. Beneath blue horizontal line, the cup tilt is too small to allow dice to escape a cup that does not extend through the table. f) Stacking measures over time for three trials, two stacking and one non-stacking. In both stacking trials, *α* and PV1 finish at 0^*°*^ and 100%, respectively. In one of the stacking trials, VR1_close_ is also 1, indicating that the dice are flush against each other.

Given that humans can perform dice stacking, one might expect it to rely on feedback-based control, as in other tasks like walking or running [4, 5]. Yet, several clues suggest otherwise. Naive participants can learn the skill within minutes, it can be done blindfolded, and tall stacks with tens of dice are possible. People likely cannot infer so many dice states through haptic feedback. We therefore hypothesize that dice stacking can be achieved, at least partly, without dice-state feedback. Here, using two simulation frameworks and a physical robot, we find simple, open-loop cup motions that robustly stack dice from random initial conditions.

## 2 Methods

### Overview

We tested open-loop cup movements using a cylindrical cup mounted to a simplified arm (Figure 1b) with two degrees-of-freedom, namely, shoulder rotation and wrist pronation-supination. The cup attaches to the wrist with a fixed outward radial tilt *θ* (Figure 1b). We simulated these manipulations using a penalty contact model [6], a MuJoCo physics engine model [7], and physical tests with a 3D printed two-DOF robot. Cup, arm, and dice dimensions reflected realistic anthropometry and common dice sizes.

We evaluated how *N*_dice_ = 4 dice responded to sinusoidal oscillations of the arm joints. Each trial began with dice dropped into the stationary cup to settle into a random configuration, followed by a fixed duration of arm, wrist, and cup oscillations. We then evaluated dice stacked-ness. We varied wrist and shoulder oscillation amplitudes (*A*_wrist_, *A*_arm_), static wrist tilt (*θ*), and shared oscillation frequency (*ω*); see Supplementary Table S1.

### Rigid body simulations with penalty contact dynamics

We modeled the cup as a cylinder with a circular end-cap, the dice as cubes, and the table as flat and horizontal (Figure 1a). All bodies were nominally rigid, with contact forces computed from rigid body states. To simplify cup-table interaction, the cup wall extended through the table, preventing dice escape through the bottom. Normal contact forces followed simple penalty rules [6, 8], which assign forces via a linear function of the local penetration and relative speed between the interfering bodies as if springs and dampers resisted interpenetration. Tangential forces used an approximate dry friction model with a sigmoidal force-velocity curve, producing normal-force-proportional high-viscosity damping at low speeds and standard rate-independent friction at higher speeds [9]. This approximates static friction over the short dice stacking time scales. Dice-cup and dice-table contact forces acted at dice vertices, with contact normals obtained from the penetrated cup or table surface. For dice-dice interactions, forces were computed at mesh points embedded in each face, with local normals matching face normals, making the contact interactions resemble hydrostatic pressure distributions [8]. Contact parameters were minimally tuned: stiff enough to avoid large penetrations but not too stiff for computation. Dynamics were integrated using 4th order Runge-Kutta at 20 kHz (MATLAB implementation in *Supplementary Code*). We ran two sets of simulations: first, a coarse sampling of arm oscillation parameters, then a finer sampling (Table S1).

### MuJoCo model simulations

MuJoCo (Multi-Joint dynamics with Contact) is a general-purpose physics engine widely used in robotics and biomechanics [7]. MuJoCo computes rigid body contact forces via convex optimization at each step [7], minimizing a quadratic function of the post-contact velocity under friction, unilateral normal force, and non-penetration constraints. Since MuJoCo requires convex geometries, we modeled the cup wall as a union of cylindrical prisms arranged in a circle to approximate the curved inner surface (Figure 2a), and capped both ends to contain the dice. We tested the same parameter combinations as the fine sampling in our penalty contact model (Table S1), using 4th order Runge-Kutta integration at 20 kHz.

**Figure 2:**
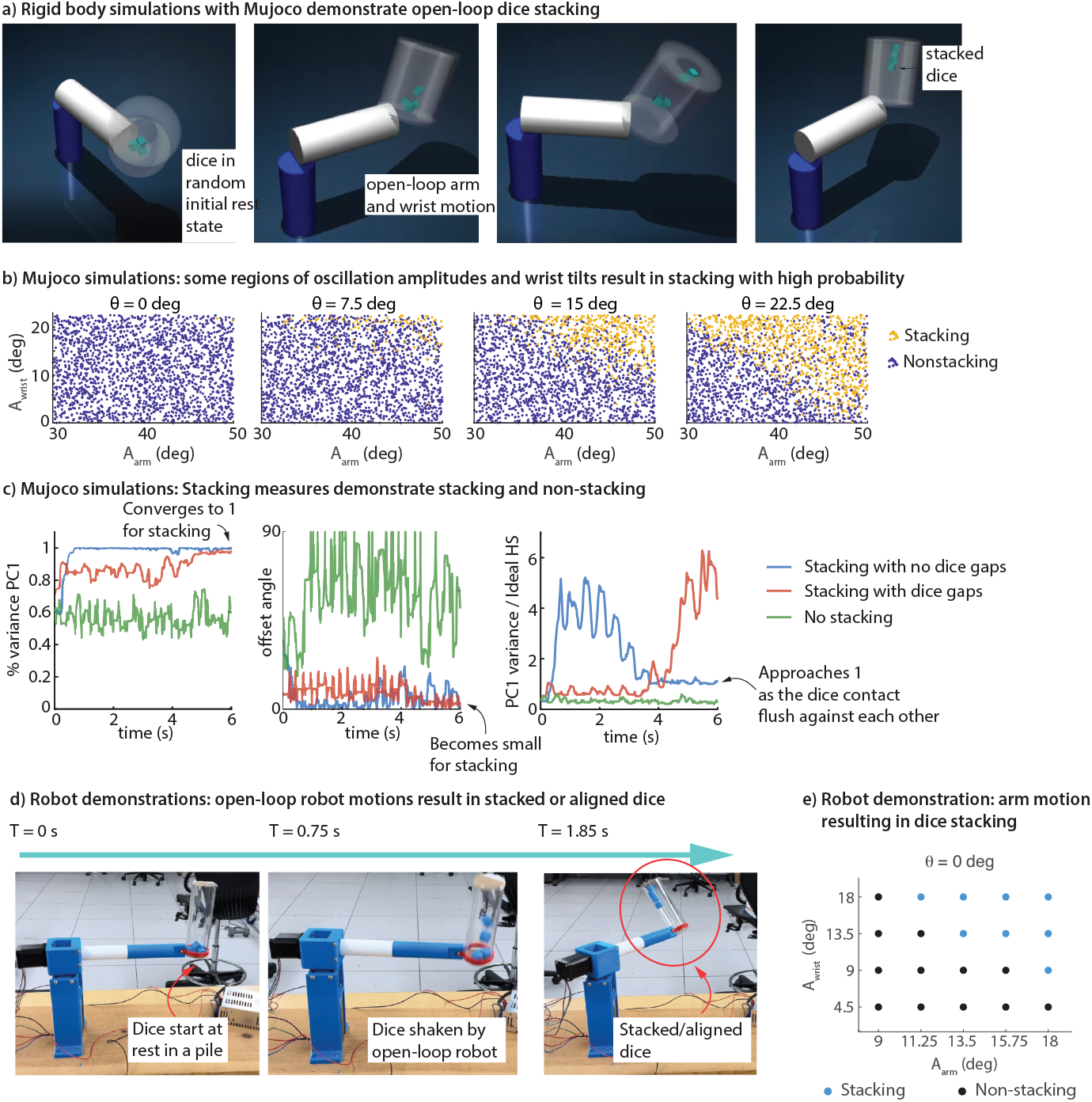
Mujoco simulation and robot demonstration exhibit open-loop dice stacking. a) Successful Mujoco stacking trial, showing the dice go from randomly arranged to being in a line and flush against the cup wall. b) Oscillation amplitutes and radial wrist tilt that result in high likelihood of stacking in mujoco simulations. c) Stacking measures for three Mujoco trials: one non-stacking and two stacking, one with flush dice and another gaps. d) 3D-printed robot demonstrating stacking from a random initial pile. e) Robot arm and wrist oscillation parameters showing clear stacking versus no-stacking regions.

### Physical robot experiments

For physical validation of these simulations, we built a two-degree-of-freedom robot to reproduce a subset of cup trajectories (Figure 2d). The robot had a capped acrylic cup with zero radial tilt (*θ* = 0). Most components were 3D-printed in PETG, with commercial actuators, bearings, and shaft couplings. Stepper motors driven by an Arduino UNO controlled the wrist and arm. We tested a small rectangular grid of arm oscillation parameters (Table S1). Dice began in random static equilibrium, followed by a fixed-duration oscillation.

### Quantifying successful stacking

To assess stacking in each computer simulation trial (Figure 1c), we used two measures based on the dice centers of mass distribution: (1) percent variance explained by the first principal component (PV1), and (2) the absolute angle between that component and the cup’s long axis (*α*). When PV1 is near one, the dice are self-aligned; when *α* is near zero, they are cup-aligned. Dice flushed against the inner wall or spaced along a line parallel to the cup’s axis therefore have PV1 near one and *α* near zero. We considered such configurations “stacked,” as they could likely form a free-standing stack after a stopping maneuver involving slowing the oscillation and reducing the static radial tilt. A third measure, VR1, which is the ratio of the observed first PC variance to that of a perfectly flush stack, captured how flush the dice were with one an other. Robot trials, lacking kinematic sensors, were judged visually. Successful stacking meant a persistent vertical column with possible small gaps.

## 3 Results

### Both computational models and the physical robot show open-loop dice stacking

Initial coarse parameter exploration (Table S1) using the penalty-contact model identified 2 Hz as the best stacking frequency. Finer exploration revealed a contiguous region in the arm–wrist oscillation amplitude space where stacking reliably occurs (Figure 1d-f). Some simulations yielded flush dice contact (VR1 *≈* 1), while others produced dice stacked with gaps (VR1 > 1)(Figure 1f).

Mujoco simulations (Figure 2a-c) showed a similar region of oscillation parameters that result in stacking, though the transition from non-stacking to stacking occurred gradually, unlike the sharper boundary in the penalty-contact model. Some stacking trials produced flush dice, others did not.

Robot experiments (Figure 2d-e) qualitatively matched both models in displaying a robust stacking region, but with a sharp boundary like the penalty-contact case.

Together, these results suggest that dice stacking, as approximated here, is at least partially open-loop: that is, the dice alignment within the moving cup can be achieved with simple motions without sensing the dice state.

## 4 Discussion

We have shown that approximations of dice stacking can be achieved via a wide range of humanly achievable, open-loop arm and cup movements. This is surprising, as the task seems impossible, let alone achievable with a dice-agnostic motion. An unpublished 2D simulation study [10] failed to learn dice stacking via reinforcement learning, so the authors learned a feedback strategy from human demonstrations. Counterintuitively, our results suggest that had they studied the task in 3D instead of 2D, their reinforcement learning attempts might have succeeded.

### A potential mechanism

A possible mechanism is suggested by an easier task: aligning dice within a cup lying horizontally on its side. Simply rolling the cup—back and forth or continuously—empirically aligns most initial dice configurations. Our dice stacking cup motions (Table S1) generate qualitatively similar dice accelerations, with centripetal acceleration during arm swings acting like gravity in the rolling case, suggesting a shared explanation. In the horizontal rolling task, alignment may arise from potential energy reduction: the lowest-energy configuration is dice aligned with the cup axis. As the cup rolls, dice reshuffle through dissipative interactions, gradually settling into lower-energy states, with rolling providing energy to escape local minima. Future work could formalize this proposed mechanism mathematically.

### Stopping maneuvers

We showed stack formation inside the oscillating cup but did not examine stopping maneuvers that end with a stationary, free-standing stack. Whether such stopping is possible with open-loop strategies remains unclear. In our practice with real dice, stopping without the stack collapsing was harder than forming or maintaining stacks in a moving cup. Because the required duration of shaking varied greatly between attempts, timing the stop depended on auditory cues. Sliding sounds suggested stacking completion, while clacking indicated more shaking was needed. Future work can examine the feasibility of various simple open-loop or closed-loop strategies for successful stopping maneuvers.

### Geometric simplifications

We ignored certain interactions between the dice, cup rim, and table. Our cup either extended through the table or had a capped bottom, preventing dice escape, unlike a real cup. However, since cup accelerations always directed dice away from potential escape regions, open cups would likely behave similarly. Future work should test this with a robot sweeping an unsealed cup across a table. Another simplification may explain some differences across frameworks: smooth-walled cups in penalty-contact and robot simulations showed a sharp stacking boundary, while the faceted Mujoco cup showed a probabilistic transition.

### A spectrum of dynamic manipulation tasks

Dynamic manipulation tasks can be viewed along a spectrum from open-loop to closed-loop control. On the open-loop side, [11] showed that thousands of dice in a bucket become ordered when the bucket oscillates above a frequency threshold. Similarly, in the Brazil nut effect [12], random shaking sorts large particles upward. On the closed-loop side lie tasks like pick-and-place [9], with greater control authority and specific goals. Between these extremes are dice stacking, some juggling [13], universal parts orienting [14], and even “percussive maintenance” (hitting an appliance to make it work). Recognizing where a task lies on this spectrum may aid learning, and perhaps humans implicitly do so. This may account for the ease with which humans learn dice stacking. In this case, reinforcement learning methods that bias toward mixed or open-loop strategies [15] could improve models of biological learning and help bridge the gap between artificial and human manipulation. Future work should test whether explicit open-loop biases are needed, as standard reinforcement learners may overfit to dice-state feedback and generalize poorly compared to open-loop strategies.

### Robustness to simulation parameters and assumptions

Observing stacking in two distinct simulations and a physical robot suggests insensitivity of the stacking to contact-model details. Moreover, the physical robot achieved stacking despite substantial kinematic variability from mechanical compliance, indicating that precise trajectories are unnecessary for successful stacking. Future work can quantify this robustness via a dynamical systems analysis (e.g., examining the fixed point basins of attractions with model assumptions) and examine whether this robustness extends to stopping maneuvers.

## Data and code accessibility

Both the MATLAB code and the python code with mujoco are available from the following webpage for review purposes: https://tinyurl.com/DiceStackingPrograms and will be available on figshare and github upon publication.

## Supplementary Information

This manuscript has a Supplementary Appendix containing Supplementary Table S1 and Supplementary Video V1, uploaded with the submission.

## Author contributions

NE and MS conceived the study. NE and ML performed the research. NE, ML, and MS participated in drafting and revising the manuscript, and all authors approved the final manuscript.

## Competing interests

We declare that the authors have no competing interests.

## Funding

NE was supported by a fellowship from the Ohio State University. Some simulations were executed on the Ohio supercomputer [16].

## Acknowledgments

The authors thank Matthew Handford for insightful early conversations that informed this work and for bringing the phenomenon to our attention.

## Supplementary Appendix for

### Supplementary Table

**Table S1:** Parameter values for arm and wrist movement used to explore stacking versus not stacking. When the ‘belongs to’ symbol ∈ is used, the parameter value is chosen from one of the finitely many values in the set shown.

| Parameter | Initial coarse sample<br>(bespoke model only) | Finer sample<br>(both models) | Robot sample |
| --- | --- | --- | --- |
| Wrist amplitude $A_{\text{wrist}}$ (degrees) | $\in \{0, 22.5\}$ | Drawn uniformly from $(0, 22.5)$ | $\in \{4.5, 9, 13.5, 18\}$ |
| Arm amplitude $A_{\text{arm}}$ (degrees) | $\in \{30, 45, 60\}$ | Drawn uniformly from $(30, 50)$ | $\in \{9, 11.25, 13.5, 15.75, 18\}$ |
| Wrist outward tilt $\theta$ (degrees) | $\in \{0, 9, 18, 27\}$ | $\in \{0, 7.5, 15, 22.5\}$ | 0 |
| Oscillation frequency $\omega$ (Hz) | $\in \{1, 15/14, \dots, 2\}$ | 2 | 1.6 |
| Initial settling duration (seconds) | 0.2 | 0.2 (bespoke) or 1 (Mujoco) | unmeasured |
| Oscillation duration (seconds) | 2 | 2 (bespoke) or 4 (Mujoco) | 10 |

## Notes

https://github.com/Lerner-Ma/dice-stacking-simulation

### Competing Interest Statement

The authors have declared no competing interest.

### Summary of Updates

Reformatted to remove any journal-specific branding; Added a supplementary table containing details about our simulation and robot experiments.

